# Identifying the potential for sustainable human–wildlife coexistence by integrating willingness to coexist with habitat suitability models

**DOI:** 10.1101/2022.09.02.506181

**Authors:** Susanne Marieke Vogel, Divya Vasudev, Joseph O. Ogutu, Purity Taek, Emilio Berti, Varun R. Goswami, Michael Kaelo, Robert Buitenwerf, Michael Munk, Wang Li, Jake Wall, Desalegn Chala Gelete, Irene Amoke, Alice Odingo, Jens-Christian Svenning

## Abstract

Persistence of large mammals in the Anthropocene depends on human willingness to coexist with them, but this is rarely incorporated into habitat suitability or conservation priority assessments. We propose a framework that integrates human willingness-to-coexist with habitat suitability assessments to identify areas of high potential for sustainable coexistence. We demonstrate its applicability for elephants and rhinos in the socio-ecological system of Maasai Mara, Kenya, by integrating spatial distributions of people’s willingness-to-coexist based on Bayesian hierarchical models using 556 household interviews, with socio-ecological habitat suitability mapping validated with long-term elephant observations from aerial surveys. Willingness-to-coexist was higher if people had little personal experience with a species, and strongly reduced by experiencing a species as a threat to humans. The sustainable coexistence potential framework highlights areas of low socio-ecological suitability, and areas that require more effort to increase positive stakeholder engagement to achieve long-term persistence of large herbivores in human-dominated landscapes.

## 1. Introduction

Human coexistence with wildlife is increasingly recognised as key to successful conservation and restoration in areas where humans and large mammals share space and resources (Frank & Glikman, 2003; König et al., 2020). But if people are unwilling to coexist—share space and resources—with large mammals, the prospect of their long-term persistence or population growth will be low in the shared spaces, even in socio-ecologically suitable habitats (Franco and Eivin, 2021; Pebsworth and Radhakrishna, 2021). This can be linked to intensive human–wildlife and human–human conflicts, to the detriment of wildlife, local communities and conservation programs (Dickman, 2010). Ignoring willingness to coexist when assessing coexistence potential (e.g. Cretois et al. 2021) thus limits the practicability of potential conservation interventions. In addition to the assessments of wildlife needs and constraints, it is therefore necessary to incorporate the perpectives of the people coexisting with wildlife into conservation research and policies (Ardiantiono et al., 2021; Bruskotter & Wilson, 2014; Frank & Glikman, 2003; St John et al., 2010).

By combined evaluation of socio-ecological spatial variables expected to be relevant for wildlife, and the habitat use of those species, habitat suitability assessments can be considered as ‘letting’ animals voice their requirements (Buller, 2015), or more specifically, a human perspective on the needs and preferences of animals (Lötter et al. 2008; Wemmer & Christen, 2008). Joining this with a measure of the potential of human-wildlife coexistence from a human perspective (König et al., 2020) by integrating it with willingness to coexist, leads to a more sustainable –and realistic– measure of the potential for human–wildlife coexistence. This results in a non-anthropocentric perspective that values both human and non-human lives (Born et al., 2001), which aims to find a common ground between people-centred conservation and science-led ecocentrism (Sandbrook et al. 2019). Furthermore, the success of spatial conservation prioritisation strategies reveals the opportunities of applying a concept of conservation priority zoning to achieve the integration of biological and social values (Whitehead et al., 2014).

Attitudes are linked to behavioural intent and behaviour, and can be considered to follow a scale from active negative resistance, to passive tolerance, and finally active positive stewardship (Bruskotter and Fulton, 2012; Treves, 2012). There is however a bias towards research on negative experiences and attitude towards unwanted species instead of their acceptance, which may tend to oversimplify our understanding of the plurality of people’s perspectives and experiences with wildlife (Bruskotter and Wilson, 2014; Buijs and Jacobs, 2021). Since attitudinal research relies on self-reporting, there is also a risk of false positive and false negative reporting errors, as people can overstate or understate their opinions (sub-) consciously (Vasudev et al., 2020; Vasudev and Goswami, 2019).

Evidence for a clear link between attitude and actual behaviour of people is currently limited (Ajzen and Fishbein, 2021; Liu et al., 2011; St John et al., 2010). For example, there is limited quantitative evidence for a link between stakeholder attitudes and the socio-ecological suitability of an area for large herbivores such as elephants (*Loxodonta africana*) and black rhinoceros (*Diceros bicornis;* Vogel et al., 2021). Nevertheless, a positive attitude can be linked to behaviour supporting conservation at a local scale, and can thus be expected to increase suitability of an area as a habitat or movement conduit for a species (Behr et al. 2017; Broekhuis et al., 2018; Liu et al., 2011; Vasudev et al., 2020). To successfully integrate anthropogenic and ecological perspectives into holistic habitat assessments, we need to recognize and overcome these challenges of negative research bias, self-reporting errors and limited established links between attitude and conservation success.

To incorporate attitudes in habitat suitability assessments, we therefore calculated a ‘willingness-to- coexist’ score which represents *the probability that people have a positive attitude towards sharing space and resources with wildlife on a local scale*. We do so using hierarchical Bayesian models that account for misreporting errors, developed by Vasudev & Goswami (2019) based on the occupancy modelling framework used in ecology. While anyone can have a notional, or generic attitude toward animals, willingness-to-coexist (or its lack thereof) follows from, and pertains to, living in close contact with wildlife on a local scale (Vasudev et al., 2020; Vasudev and Goswami, 2019). We consider willingness-to-coexist to be a higher-order or active positive attitude, meaning that it is closer to active support and stewardship than normative attitudes, basic beliefs and values (Bruskotter and Fulton, 2012; Fulton et al., 1996). Willingness-to-coexist thus goes beyond mere inaction (Treves, 2012) or passive tolerance (Bruskotter & Wilson, 2014), addressing the aforementioned biases in attitudinal assessments.

We integrate willingness-to-coexist scores with potential elephant densities based on socio- ecological suitability of areas, calculated using habitat suitability models. The result is a tool that can aid spatial planning for coexistence in a human-dominated world by highlighting areas of low willingness to coexist or low habitat suitability, and identify conservation priorities (Broekhuis et al., 2017; Marchini et al., 2019). From a wildlife persistence perspective, there is a socio-ecological threshold below which the potential for wildlife occurrence is too low for the landscape to fulfil the species’ needs (Figure 1a; Svenning & Faurby 2017). Yet, to achieve sustainable coexistence, both animal and human occurrence potential thresholds must be met (Jochum et al., 2014; Figure 1b), with sufficiently suitable environmental conditions and the satisfaction of human interests *without depriving species, native ecosystems or native populations of their health’* (Vucetich & Nelson 2010). Without the later, continued human support for wildlife is unlikely (Franco and Eivin, 2021; Pebsworth and Radhakrishna, 2021).

**Figure 1.**
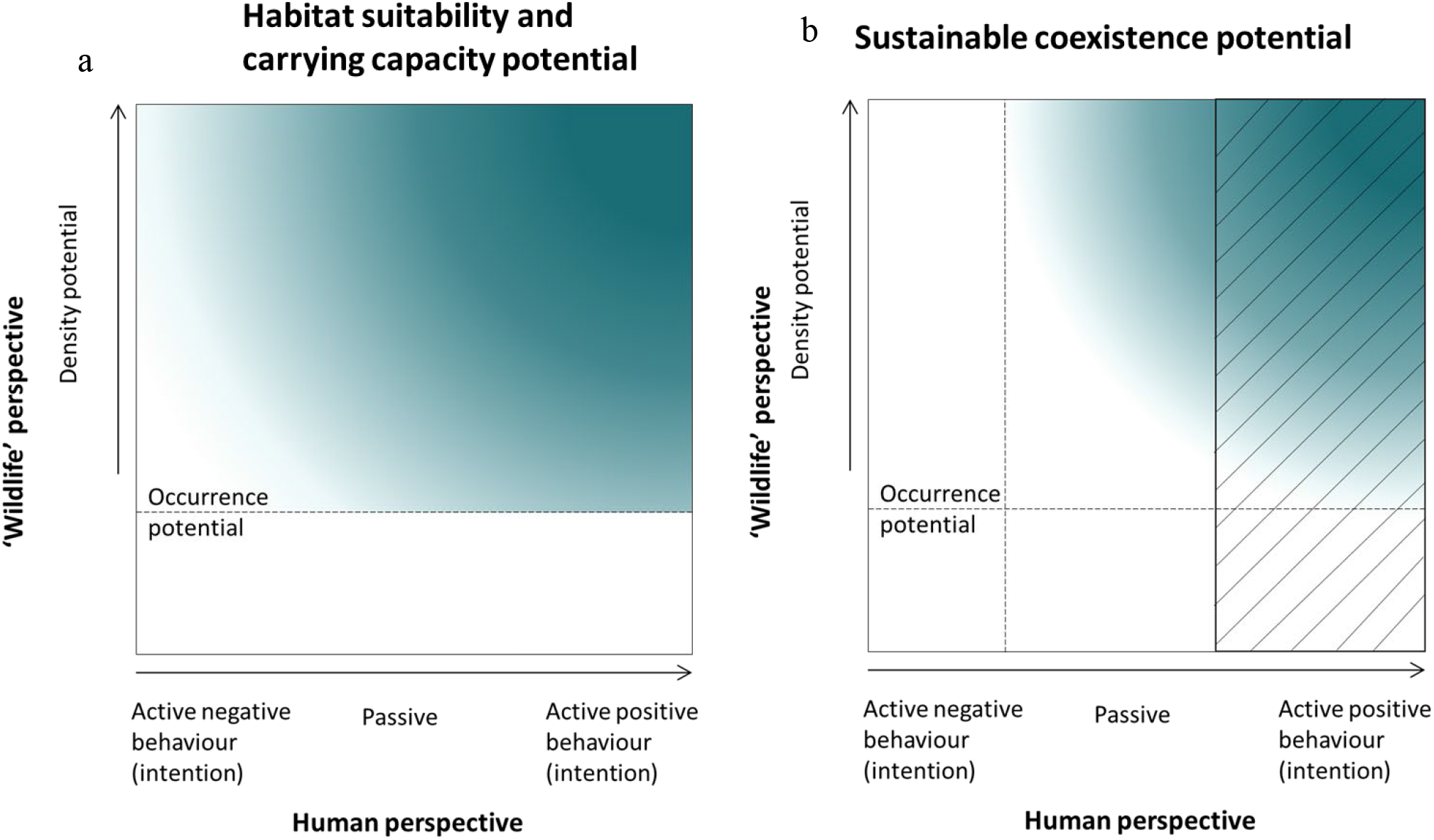
Conceptual diagram representing. **a)** socio-ecological habitat suitability for wildlife (occurrence potential) and carrying capacity (density potential), and **b)** sustainable human wildlife coexistence potential. The x-axes represent a negative-positive scale for attitude, behavioural intent and behaviour of people, and the y-axes represent wildlife density potential. In **a)**, we assume that there is some level of potential for occurrence of wildlife across the anthropogenic scale and that an ecological threshold exists below which wildlife cannot occur. In **b)** we assume a threshold exists for both axes below which sustainable coexistence is not possible. The darker the colour, the higher is the suitability and carrying capacity potential **(a)** or potential for sustainable coexistence **(b)**. The hatched area in b) represents the space where people can be expected to express a willingness-to-coexist, an attitudinal measure for positive active stewardship.

Besides sufficient resources, this requires socially just conservation through effective institutions that aim to satisfy the needs and desires of both people and wildlife, while minimizing interactions that jeopardise each other’s safety, and result in a sustainable dynamic state in which people and wildlife co-adapt, and thus coexist (Büscher and Fletcher, 2019; Carter and Linnell, 2016; Frank and Glikman, 2019; Vucetich et al., 2018). While passiveness may be sufficient to achieve wildlife conservation, less active negative behaviour (e.g. political acts such as voting or signing petitions) could still negatively affect wildlife’s ability to survive or thrive in the long-term (Bruskotter and Fulton, 2012). Therefore, willingness-to-coexist corresponds to active positive behavioural intention (Figure 1b, hatched area). Sustainable coexistence potential should be the highest when there is potential for high densities of wildlife based on socio-ecological suitability of the landscape, and high levels of peoples’ expressed willingness-to-coexist (Figure 1b, upper right corner).

We apply the framework of sustainable coexistence potential to African savannah elephants and black rhinos in the Maasai Mara Ecosystem, Kenya. In this ecosystem there are concerns that consequences of human activities and livestock numbers are compromising wildlife persistence and movement connectivity (Ogutu et al., 2016, 2011; Stabach et al., 2022). There is also a long history of land appropriation for conservation, disempowering the pastoralist Maasai communities, weakening their traditional coping strategies, and deepening their livelihood vulnerability, thereby creating disincentives for conservation (Fernández-Llamazares et al. 2020; Goldman, 2011). Coexistence between people and African savannah elephants is of high conservation interest and concern given the elephant’s currently endangered status (Knox et al., 2021), and has significant implications for the wellbeing of people living in conservancies and pastoral lands around the Maasai Mara National Reserve (Nyumba et al., 2020).

## 2. Methods

### 2.1 Study Site

Our study site included the Naboisho, Ol Kinyei, Mara North, Lemek, Ol Chorro Oiroua, Olare-Motorogi, Proposed Muntoroben Conservancy, Olarro, Siana Mara and Olderkesi conservancies, Nyakweri Forest Conservation Area and the proposed Pardamat Conservation Area which we divided into the northern, middle and southern sections (Figure 2). We also included Aitong, Talek and Mara Rianta, three of the largest towns in the ecosystem (MMWCA, 2019).

**Figure 2.**
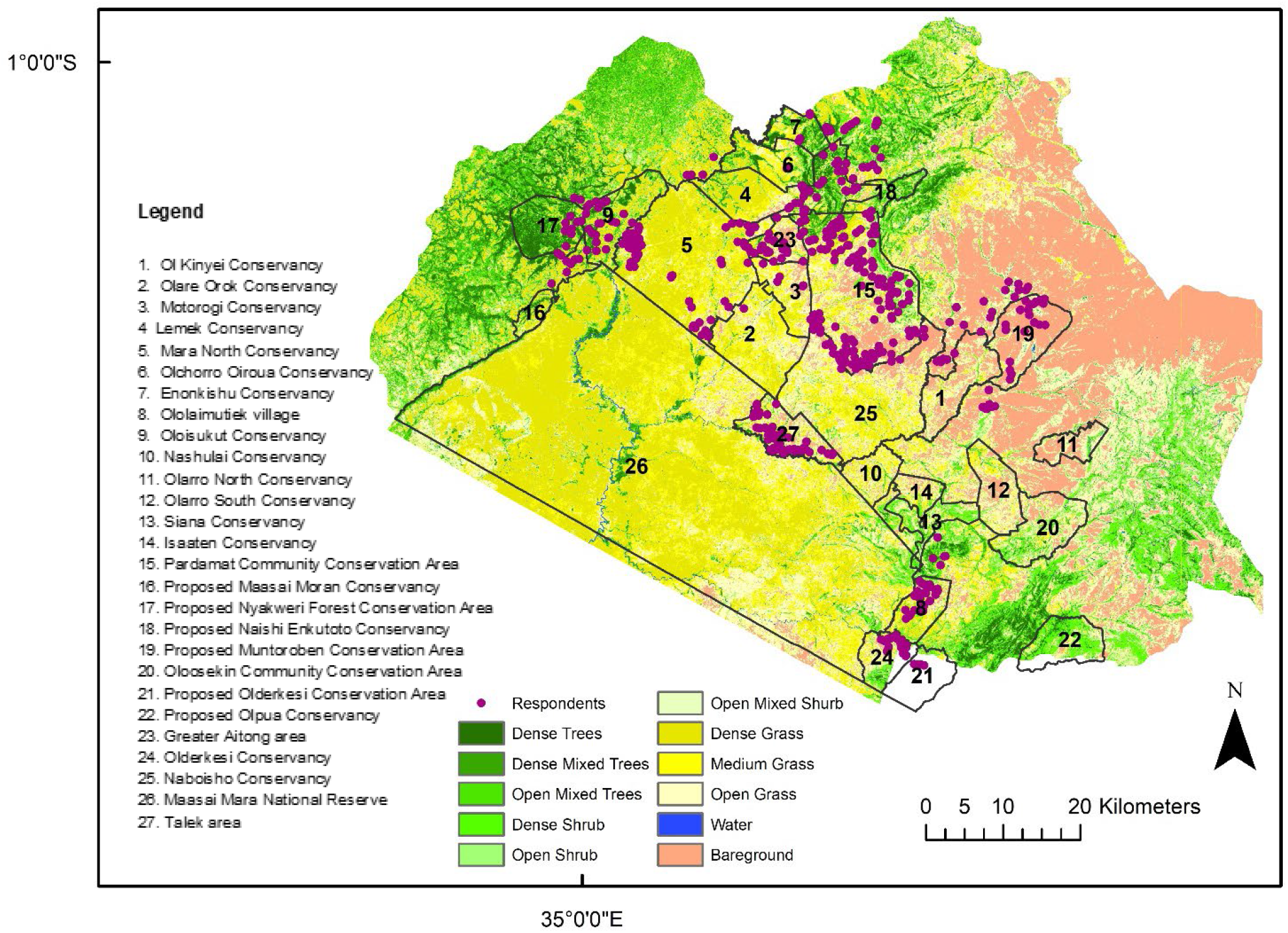
Location of respondents interviewed during the study, the names and demarcations of Maasai Mara National Reserve, (proposed) conservation areas, villages, towns and community lands and most dominant vegetation types across the study site. Most of the study site is covered by dense to medium-dense grasslands and open shrubland, interspersed with clusters of trees and grasses typically fringing drainage lines and on ridges, and bareground around areas such as in Talek and Aitong towns, and in the north-east of the study site

Land around the Maasai Mara National Reserve in Kenya is either communally or individually owned. Some landowners have organised themselves into conservancies aimed at community-based conservation, as part of the Maasai Mara Wildlife Conservancies Association (MMWCA), through partnerships with investors in eco-tourism enterprises and receive direct financial benefits, e.g., from land leases (MMWCA, 2019; Thompson et al. 2009). Elephants (3404 ± 1156 animals or 0.54 animals /km^2^ in 2022) are common throughout the ecosystem but black rhinos are rare (< 40 animals or < 0.006 animals/km^2^) and restricted to the Maasai Mara National Reserve (MMNR), and are actively pushed back by rangers into the MMNR whenever they wander into the adjoining areas to reduce the likelihood of negative human-rhino interactions. Therefore, we focused on both elephants and rhinos in the attitude assessments, but only on elephants in the potential habitat suitability and sustainable coexistence assessments.

### 2.2 Willingness to coexist model

#### 2.2.1 Interviews

We collected interview data during January - April 2020 in Maa. Interviewers were 10 men and one woman, all Maasai. Respondents were selected through stratified random sampling of 60 households from each trading centre and (proposed) conservancy, using the most complete spatial dataset on household distribution available for this region (Broekhuis et al., 2017). A total of 556 households were interviewed with each interviewer covering 40–60 households, with a non-response rate of <1%. Each interview took around 30 minutes to complete on average.

During the interviews we collected social and demographic information, asked about preferred elephant and rhino densities, experiences with both species, and collected responses to both notional and localized attitudinal statements on a five point Likert scale (Bernard, 2006; Vasudev & Goswami, 2019); notional attitudinal statements were abstract, while localized statements related directly to the respondent’s lives and livelihoods. We used the free Cybertracker software to develop an English and a Maa (most commonly spoken language among Maasai) data collection application for each of the sites and to upload these on to the interviewers’ smartphones. We also developed a measurement of acceptable wildlife density levels in areas of low literacy rates to interpret our results.

#### 2.2.2 Model construction and evaluation

To measure the probability of a positive attitude towards living with elephants and rhinos —i.e. willingness-to-coexist--we grouped the negative and passive responses to attitudinal questions into non-positive attitudes and compared these to positive responses. We applied a Bayesian hierarchical model related to ecological multiple-detection models (Miller et al. 2011; Vasudev & Goswami, 2020), which allowed us to determine false positive and false negative errors (misreporting errors) in reported attitudes, and therefore provides a more reliable measure of acceptance than methods based on self-reporting only (Vasudev and Goswami, 2019). To do so, our survey included uncertain questions, where we accounted for potential false positive and false negative response from respondents, and certain questions to which we reasonably expected only either a false positive or a false negative response. The model enables quantifying both a notional or generic attitude (ψ) and a localized attitude (φ) towards elephants and rhinos, while accounting for reporting biases (Vasudev et al., 2020; Vasudev and Goswami, 2019).

We ran three models (Model I–III), each with different sets of variables. As social-demographic information from respondents we included age, gender, the highest level of formal education, owning livestock and having an alternative livelihood strategy as covariates in all our models. Model I also included a binary variable indicating whether respondents lived inside a conservancy, as we expected this to play an important role in explaining willingness-to-coexist. We also included spatially explicit variables on reported crop loss, property loss, threat to humans, threat to livestock, and benefit from tourism. It was not possible however to include these in the other two models due to high spatial correlation between the variables. Model II included, instead of the binary conservancy/non-conservancy variable, the more spatially explicit covariate indicating the identity of each of the (proposed) conservancies and trading centres (towns). However, some conservancies contained few respondents, making results from this model less reliable. Therefore, we also fitted Model III in which respondents were divided into 11 spatial clusters with similar numbers of respondents based on spatial proximity and (potential) conservancy or trading centre. In this manuscript we focus on the results from Model I and III.

As we constructed the Likert scale by averaging at least eight Likert items on a five-point scale each, we consider it meaningful to compare means between the spatial clusters (Norman, 2010; Wu and Leung, 2017), to illustrate the deleterious effect and thus the importance of, correcting for reporting errors (Barry, 2017).

### 2.3 Habitat Suitability Models

#### 2.3.1 Species occurrence and density data

We used decadal averages of elephant numbers obtained from aerial survey monitoring data by the Directorate of Resource Surveys and Remote Sensing of Kenya (DRSRS, Ogutu et al. 2016). The decadal averages were computed for 284 sampling units, each of size 5 × 5 km and located in the MMNR as well as most of the conservancies. Surveys were separately conducted for the wet season of 2015, 2016 (two surveys), 2018 and 2022, and dry season of 2010, 2011, 2012, 2013 and 2014, and adjusted for the sampling fraction used in each survey, with sampling fraction varying from 5- 11%. Our main purpose with this particular habitat suitability model was to illustrate the potential of integrating the willingness-to-coexist mapping with habitat suitability maps. Most of the attitudinal survey data was collected during a prolonged dry season, a period during which coexistence challenges are most pronounced in the study site (Bedelian & Ogutu, 2017; Miller et al. 2014). Thus, we focus on the dry season survey data (which comprised most of our elephant survey) for mapping.

#### 2.3.2 Spatial data and covariate selection

We identified relevant ecological and anthropogenic predictors of elephant presence and density in an extensive literature review of factors known to influence elephant and rhino habitat suitability (Vogel et al., 2021) and based our final set of covariates included in the species distribution model on (multi-) collinearity between identified relevant covariates. Based on these assessments, the final covariates we included in our habitat suitability models were the proportion of the 5 × 5 km grid cells with permanent water, average terrain elevation, average terrain slope, average NDVI, livestock biomass (kg) and total number of houses in each 5 × 5 km grid cell (i.e. settlement density).

#### 2.3.3 Zero-inflated Habitat Suitability Models

The aerial surveys provided us with presence-absence data, yet the method of data collection and zero-inflation tests indicated the data was zero-inflated (Martin et al., 2005; Zuur et al. 2009; Lüdecke et al. 2021). We therefore applied two types of habitat suitability models that take zero-inflation into account, namely zero-inflated species distribution models (Poisson and negative binomial models), and spatial Integrated Nested Laplace Approximation, as described below. We constructed both types of models separately for the wet and dry season data, and both included the covariates described in section 2.3.2. As the Species Distribution Model validation also contained data from both the wet and dry seasons we also the average precipitation for each season.

#### 2.3.4 Species Distribution Model construction and evaluation

First, we used Species Distribution Models (Zero-inflated Poisson models), which are commonly used to model habitat suitability (Guisan et al., 2017). However, visualisations of the presence-absence data overlaid on the spatial covariate data layers suggested the potential presence of both sampling and structural zeros in our dataset. Consequently, we used a zero-altered negative binomial hurdle model (ZANB) to account for excess zeros taking both zero structures that emerge from unsuitable areas, and potential overdispersal (emerging from clumped distribution of elephants) into account (Blasco-Moreno et al. 2019; Martin et al., 2005; Zuur, et al. 2009) using the pscl R-package (Hadfield, 2021).

#### 2.3.5 INLA model construction and evaluation

We also modelled habitat suitability with a spatial Integrated Nested Laplace Approximation (INLA) framework that allowed us to predict elephant density through approximate Bayesian inference assuming latent Gaussian models (Moraga, 2020). We constructed the INLA habitat suitability models by using the R-INLA package (Martins et al. 2013; Rue et al. 2009; www.rinla.org). We used the zero-inflation Poisson Type 1 function because our data contains both structured and unstructured zeros (Blangiardo, 2013) and constructed them as a Besag-York-Mollié (BYM) model (Besag et al., 1991). This BYM model includes two types of random effects, an unstructured random effect that accounts for uncorrelated noise, and a Conditional Autoregressive (CAR) distribution that accounts for the spatial structure in the mapped data, by feeding into the model a neighbourhood matrix that identifies areas as neighbours if they share a boundary, and uses this information to smooth the data (Blangiardo, 2013; Moraga, 2020). We modelled both these hyper parameters using a Gaussian prior distribution *N*~ (0,1) and evaluated model fit using leave-one-out cross-validation (Blangiardo, 2013).

### 2.4 Sustainable Coexistence Potential Map

To obtain a map of Sustainable Coexistence Potential, we changed the projection and resolution of the willingness-to-coexist map (coexistence potential from the human perspective) to match the habitat suitability maps (coexistence potential from the ‘wildlife’ perspective, Figure 1). Based on frequency plots of the predicted elephant densities and willingness-to-coexist scores, we reclassified both habitat suitability maps and the willingness-to-coexist map to convert them into a binary map. We then created a combined mosaic raster plot overlapping the binary maps and summing the values of both maps, and clipped this to the area for which we had elephant density estimates. We also overlaid a difference map to highlight the areas which were socio-ecologically suitable, but not so from a human perspective, and vice versa.

### 2.5 Authors reflexivity statement and permits

The authors of this paper come from a diversity of geographical backgrounds because one of our goal was to increase knowledge exchange and address previously identified lack of collaborations within the field of elephant and rhino conservation (Vogel et al., 2021). We also have different levels and types of academic experience, different perceptions on conservation and human wildlife conflict and coexistence, and come from different ecological and social science fields. Nevertheless, the majority of the authors believe that maintaining wild large herbivores has both intrinsic and ecological values. While we aimed to learn from the Maasai perspective on living with such animals, and to incorporate the lessons learnt into common habitat suitability assessments, we do not consider this study a suitable platform for sharing in-depth Maasai perspectives on living with wildlife or revealing their issues with wildlife or conservation agents. Rather, we view this study as only one contribution to understanding (sustainable) human-wildlife coexistence in the area, highlighting the importance of integrating ecological and human perspectives when assessing the potential for coexistence and suggesting ways to do so. This research has been reviewed and approved in accordance with Aarhus University’s guidelines for the university’s Research Ethics Committee (IRB), and was completed under NACOSTI permit License No: NACOSTI/P/19/3156.

## 3. Results

### 3.1 Notional attitude towards elephants and rhinos

In general, people in the Maasai Mara had a more positive notional (or generic) attitude towards rhinos (*ψ*_rhmo_ = 0.91 [95% CrI = 0.82–0.96]), than towards elephants (*ψ*_elephant_ = 0.66 [95% CrI = 0.40–0.81]; **SI** Figure S4 a-b). The false negative reporting error was just over 30% for elephants (p_uc_^10^ = 0.35, **SI** Table S2 for the non-zero Credible Intervals and certain question reporting errors for each species) and rhinos (p_uc_^10^ = 0.32), while the false positive reporting error was higher for elephants than rhinos (p_uc_^01^, _elephant_ = 0.57, p_uc_^01^, _rhino_ = 0.30). This shows that a larger proportion of the respondents who were not positive towards elephants tended to report positive attitudes towards rhinos. The influences of covariates on these attitudes were consistent for both elephants and rhinos. Men had a higher likelihood of being positive towards elephants and rhinos notionally, while there was some tendency for older educated persons to be positive towards elephants and rhinos (Figure 2, **SI** Figure 4 a-b).

### 2.1 Willingness-to-coexist with elephants and rhinos

Respondents were generally more positive towards rhinos than elephants (Figure 2, **SI** Figure S4 a- b). There was a broad consensus among the respondents that both animal species are intelligent, important, have a right to live in the Mara, and that people are happy the animals are in Maasai Mara, occur in and use their conservancies and should continue living in the area. After correcting for reporting errors, willingness-to-coexist —relating to a localized positive attitude conditional on a notional positive attitude— was generally higher for rhinos (*ϖ*_rhino weighted average_ = 0.98 [95% CrI _weighted average_ = 0.93–1.00]) than for elephants (*ϖ*_elephant weighted average_ = 0. 59 [95% CrI _weighted average_ = 0.24–0.89], Figure 2, **SI** Figure S4 a-b). Willingness-to-coexist with elephants also varied strongly across the Maasai Mara, as with the probability of having a positive attitude towards living with elephants; from 0.13 for Talek town (s.d.=0.01) to 0.62 for the south of Paradamat (s.d.=0.00), close to Naboisho Conservancy (**SI** Table **S3** for all weighted willingness-to-coexist scores).

**Figure 2.**
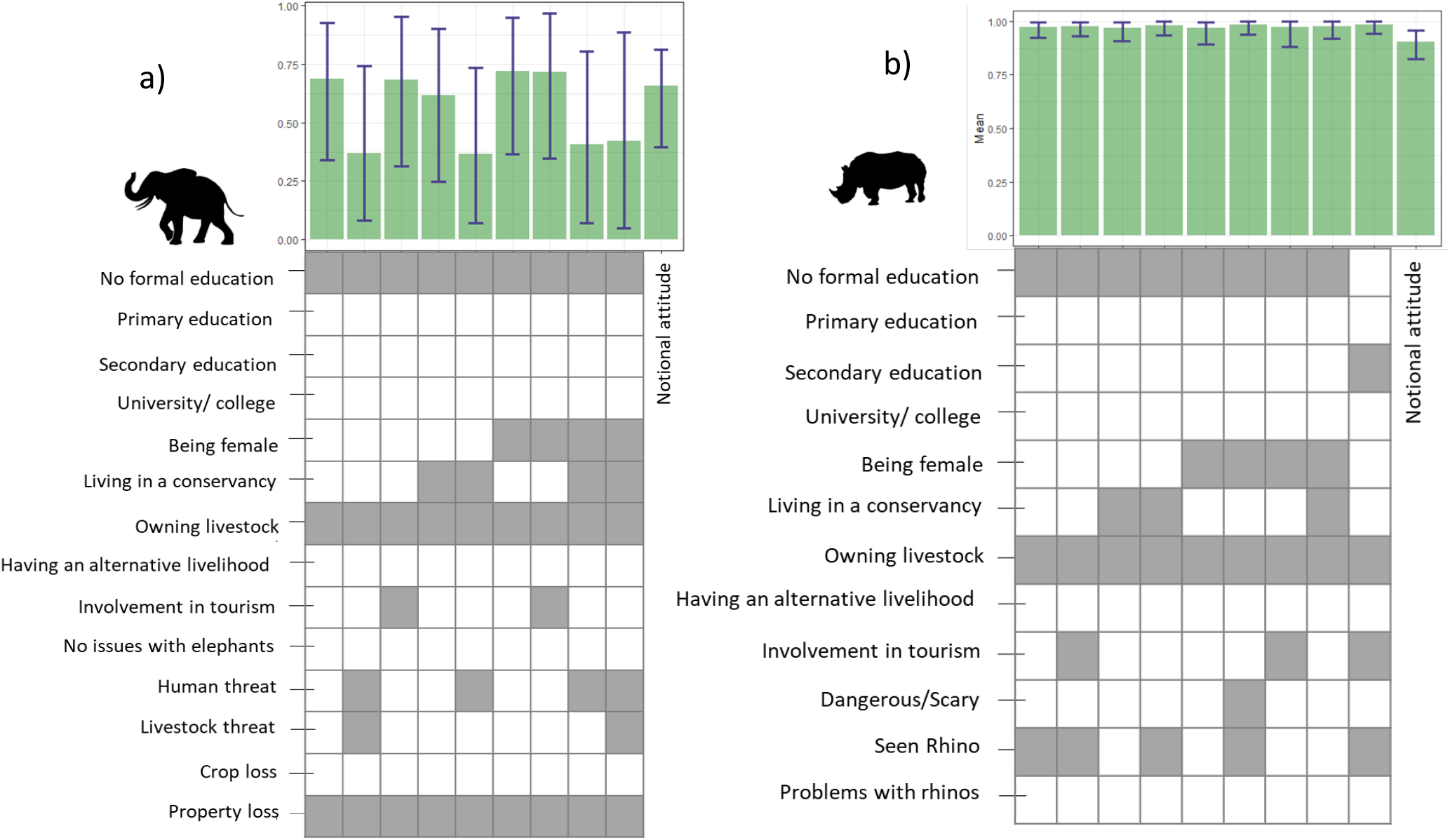
The notional attitude (ψ), and willingness-to-coexist (φ) for the most common combinations of social demographic characteristics among the respondents (n=556), expressed in the bar plot as the probability of having a positive attitude to live with elephants **(a)** and rhinos **(b)**, with 95% Credible Intervals. The x-axes of the bar plot are visualized by a matrix column representing combinations of social demographic characteristics (except for notional attitude or probability to have a positive attitude in general). For example, the first bar in Figure 2a represents the willingness-to-coexist for a man with no formal education, who owns livestock and has experienced property loss due to damage by elephants.

#### 3.1.1 Importance of correcting for reporting error

The false negative reporting error was similar for elephants (q_uc_^10^=0.42, **SI** Table **S2** for the non-zero Credible Intervals and for certain questions, the reporting errors for each species) and rhinos (q_uc_^10^= 0.41), whereas the false positive reporting error was higher for elephants than for rhinos (q_uc_^01^, _elephant_ =0.55, q_uc_^01^, _rhino_=0.42). If we would not have quantified reporting error and corrected for this in our willingness-to-coexist scores, and thus only used the Likert scores as commonly done in attitudinal studies, only support for living with elephants in Talek, the Greater Aitong area and the cluster combining Lemek, Olchoro Oiroua and Enonkishu would have been correctly estimated (**SI** Table S1). We would have overestimated support for coexistence in five, and underestimated it in four of the spatial clusters. Also, if reporting error was not corrected for when estimating the Wildlife Acceptance Capacity (WAC) indicator, then the support for coexistence with elephants would have been overestimated for three and underestimated for four of the spatial clusters (**SI** Table S1, Figure S1).

#### 3.1.2 Threat to humans

There was a stronger effect of covariates in determining people’s willingness to coexist with elephants than with rhinos, and less variance in answers among respondents (Figure 2). In particular, the threat it poses strongly reduces people’s willingness-to-coexist with elephants. Where willingness-to-coexist with them was low, elephants were perceived as posing greater threats to human life than other threats (Figure 2a; **SI** Figure S3, S4 c-d). Elephants obstructed people’s daily activities, and were described as *“causing fear and frustration in human[s] if they come around”, “They have jammed every space around here. We fear them a lot.”* Some of the open answers people gave also indicated the complexities of the issues involved, for example, *“They disturb our school going children, [but] on the other hand there are no means of transport, and the school is very far”*. At the same time, people expressed that *“these animals are bringing a lot of income into this country, therefore maximum protection is needed [for] this species*” and that “*elephants and rhinos [should] be secured for the future generations”*.

#### 3.1.3 Personal experience

Willingness-to-coexist with elephants tended to be lower among younger respondents, and higher among people with primary education than those with no formal education. Those who had seen rhinos and owned livestock were more willing to coexist with rhinos (Figure 2, **SI** Figure S4). Many respondents emphasised their limited experience with rhinos, *“I’ve never seen a live rhino. Only on phone pictures”*, and “*I saw one 15 years ago. It was an amazing experience”*. This also resulted in inconsistencies in the description of the animals, both on its appearance *“it’s like [a] hippo but the difference is [that] the rhinos have the horn”, “it’s like [a] warthog”* or people referring to it as a *“brownish elephant-like monster”;* on the animal’s character, descriptions varied from *“very rude”, “very stubborn”, “very hateful to people”*, to *“very polite, it’s a shy animal”* and a *“very friendly animal*”.

### 3.2 Habitat Suitability Model I: Species Distribution Model (SDM)

In the final selected model used to construct the potential elephant density map (Figure 5b), several ecological variables were significant in predicting both the density and occurrence probability of elephants (Table 1). Precipitation (seasonality) was an important predictor of elephant absences, but not their density. NDVI and Elevation were important as predictors of elephant density but not their absence (Table 1).

**Table 1.** Parameter estimates and the associated standard errors for the truncated negative binomial with a log link function constituting the count part of the hurdle model used to predict the potential elephant density, and the binary model with a logit link function constituting the zero part of the hurdle model used to predict the potential presence of elephants across the Maasai Mara. The Z-test tests if a parameter estimate is different from zero. Significant variables are highlighted in bold face font.

|  | Variable | Estimate | Std. Error | Z-value | P-value |
| --- | --- | --- | --- | --- | --- |
| Truncated<br>negative<br>binomial with a<br>log link<br>function | **Intercept** | **2.982** | **0.148** | **20.089** | **<0.000** |
|  | Water | 0.072 | 0.081 | 0.0886 | 0.376 |
|  | NDVI | 0.153 | 0.178 | 0.856 | 0.392 |
|  | Elevation | 0.165 | 0.150 | 1.097 | 0.273 |
|  | Total number of<br>houses | -0.087 | 0.197 | -0.444 | 0.657 |
|  | **Precipitation** | **0.312** | **0.122** | **2.614** | **<0.001** |
| Binary model<br>with a logit link<br>function | **Intercept** | **-0.699** | **0.133** | **-5.254** | **<0.000** |
|  | Water | -0.077 | 0.123 | -0.600 | 0.550 |
|  | **NDVI** | **0.623** | **0.163** | **3.860** | **<0.000** |
|  | **Elevation** | **-0.990** | **0.159** | **-6.218** | **<0.000** |
|  | Total number of<br>houses | -0.089 | 0.213 | -0.413 | 0.201 |
|  | Precipitation | 0.173 | 0.136 | 1.279 | 0.595 |

**Figure 5.**
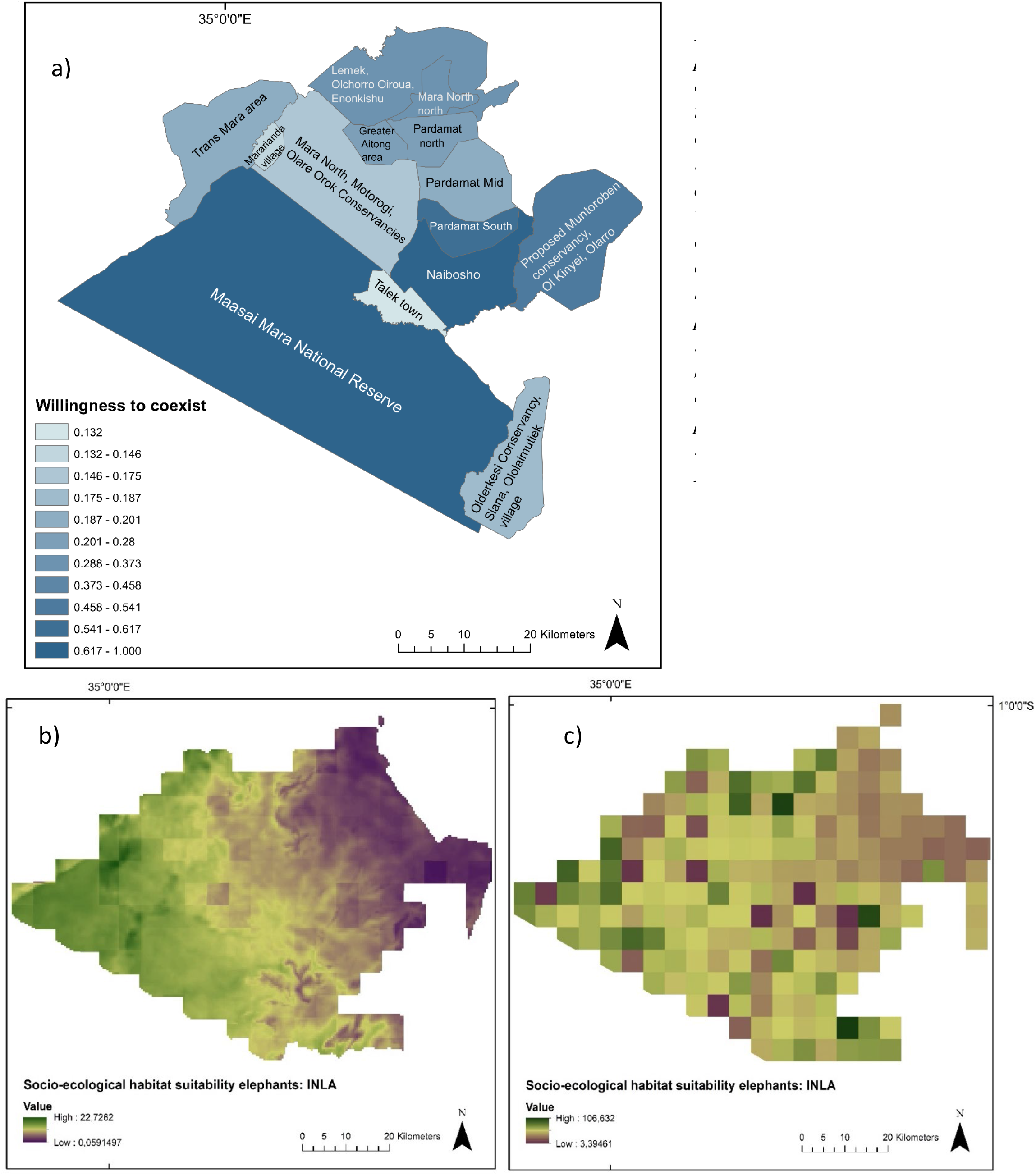
**a)** Probability of a positive attitude towards living with elephants (willingness-to-coexist) as measured across the (proposed) conservancies and other regions in the Maasai Mara, based on the average willingness-to-coexist weighted by the prevalence of a combination of social demographic characteristics (n=556); **b)** Habitat suitability map and the predicted elephant density across the Maasai Mara based on model I: SDM; **c)** Habitat suitability map and the predicted elephant density across the Maasai Mara based on model II:INLA model.

### 3.3 Habitat Suitability Model II: Integrated nested Laplace approximation (INLA)

The NDVI is positively correlated with and is the most important predictor of the potential number of elephants in the study site, based on the posterior marginal of the fixed effect parameters, calculated by smoothing the marginal distribution of the coefficients (Moraga, 2020; Table 2). Terrain slope and the amount of water present also appear to positively influence the elephant density potentials, with very small effects of elevation and the total number of houses, and no effect of livestock biomass (Table 2; Figure 5c).

**Table 2.** The estimated posterior mean and its standard deviation and quartiles for parameters of the fixed effects in the INLA model used to predict the potential elephant density across the Maasai Mara.

| *Variable* | *Mean* | *s.d.* | *0.025*<br>*quantile* | *0.500*<br>*quantile* | *0.975*<br>*quantile* |
| --- | --- | --- | --- | --- | --- |
| *Intercept* | 7.038 | 4.688 | -2.913 | 7.306 | 15.498 |
| *Water* | 1.692 | 15.218 | -28.175 | 1.702 | 31.497 |
| *NDVI* | 8.682 | 3.651 | 1.545 | 8.674 | 15.863 |
| *Elevation* | -0.006 | 0.002 | -0.010 | -0.006 | -0.001 |
| *Slope* | 0.129 | 0.074 | -0.017 | 0.129 | 0.275 |
| *Total number of*<br>*houses* | 0.013 | 0.006 | 0.000 | 0.013 | 0.024 |
| *Biomass of livestock* | 0.000 | 0.000 | 0.000 | 0.000 | 0.000 |

### 3.4 Sustainable coexistence potential

The spatial distribution of sustainable coexistence potential in the Maasai Mara conservancies is the highest around Naboisho, Pardamat South and the Proposed Muntoroben Conservancy, Ol Kinyei and Olarro conservancies as both willingness-to-coexist and potential elephant densities are high in these areas (Figure 6). These areas therefore appear to fall within the top right corner of our conceptual Figure 1b. While parts of the proposed Muntoroben Conservancy, Ol Kinyei and Olarro cluster do appear to lie in the hatched area of Figure 1b with its relatively high willingness-to- coexist, lack of measured habitat suitability places this location in the bottom-right corner, above the occurrence potential line of the conceptual diagram. Talek Town, an area in between these areas and the MMNR, however has low sustainable coexistence potential due to low values of both willingness-to-coexist and habitat suitability, placing it outside the dark blue area of our willingness-to-coexist diagram (Figure 1b).

**Figure 6.**
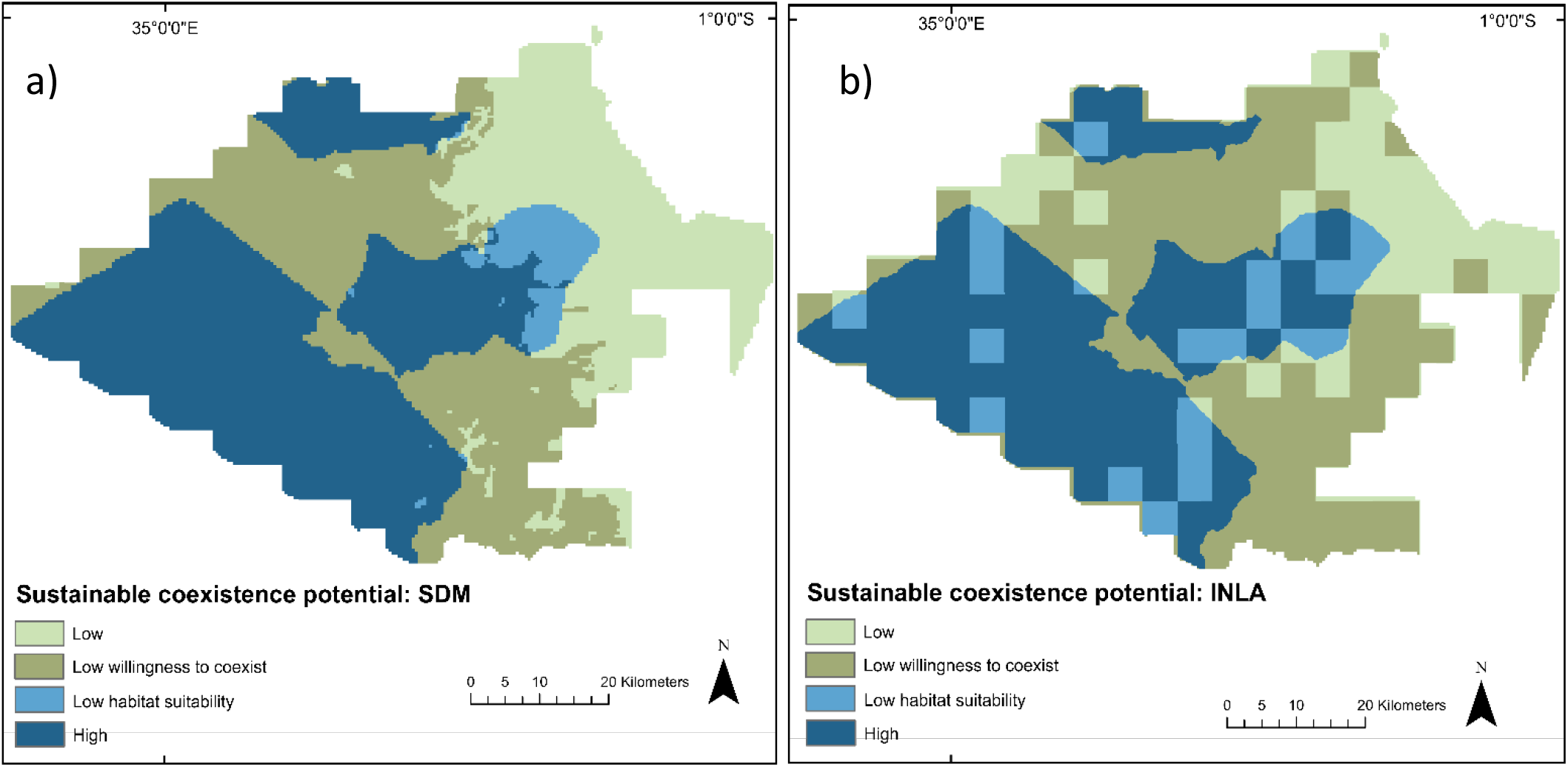
Overlaid map indicating the sustainable coexistence potential of elephants and people across the Maasai Mara, based on binary suitability as predicted by the probability of a positive attitude towards living with elephants (willingness-to-coexist) as measured across the (proposed) conservancies and other regions in the Maasai Mara. The estimated potentials are based on the average willingness-to-coexist weighted by the prevalence of a combination of social demographic characteristics (n=556) and **a)** Habitat suitability map and the predicted elephant density across the Maasai Mara based on model I: SDM; and **b)** Habitat suitability map and the predicted elephant density across the Maasai Mara based on model II:INLA model.

## 4. Discussion

We proposed and demonstrated how to integrate coexistence potential from both human (willingness-to-coexist) and wildlife (socio-ecological habitat suitability) perspectives into sustainable coexistence potential, by accounting for issues that have previously prevented the inclusion of willingness-to-coexist into habitat assessment models.

Results of our hierarchical attitudinal model highlight the impact of (negative) experiences with wildlife on people’s willingness-to-coexist. Compared to elephants, willingness-to-coexist with rhinos was higher and closer to the notional attitudes toward rhinos. The threat to their lives people perceived from living with elephants was the main factor reducing willingness-to-coexist, which is consistent with results reported for Asian elephants (Ardiantiono et al., 2021). This suggests that perceived threats to human life (Dickman, 2010; Zimmermann et al., 2020), can potentially serve as a proxy for willingness-to-coexist in spatial planning, and that targeted conservation interventions addressing this perception of risk can enhance overall potential for sustainable coexistence.

However, in other socio-ecological landscapes economic losses due to crop or livestock depredation and property damage could be more important, as except for Trans Mara, planting of crops was limited in our study site.

The results from the habitat suitability models highlight the importance of using a holistic perspective, and considering as direct proxies as possible for both the ecological and anthropogenic factors in species distribution models (Vogel et al., 2021). It also highlights the complexity and spatial variability inherent in the factors driving both elephant presence and density. While the absence and density of elephants was mainly determined by ecological variables, models performed better if anthropogenic variables were included, in particular the number of houses. The presence and density of human settlements has been previously demonstrated to have a strong negative influence on elephant space use and movement in Asia (Goswami et al., 2021, 2014). The importance of precipitation, and the NDVI, a measure of vegetation greenness, in driving elephant density predictions, suggest that wildlife in this socio-ecological system is strongly depended on rainfall, similar to people and their livestock (Miller et al., 2014). This also highlights the danger of climate change, and in particular droughts, to further reduce human–wildlife coexistence potential and increase the risks of land use change and physical conflicts between people and wildlife.

The distribution of sustainable coexistence potential for elephants and people across the Maasai Mara suggests that sustainable coexistence is contingent upon increasing habitat suitability for wildlife (Ogutu et al., 2019) without (further) stigmatizing the pastoralist community and with incorporating their experiences and willingness to share space and resources with wildlife (Cavanagh et al., 2020; Weldemichel, 2020; Weldemichel et al. 2019). We identified areas that had low habitat suitability for elephants during the dry season, but high willingness-to-coexist, highlighting an underutilized potential of such areas for promoting wildlife-human co-existence. On the other hand, we also found ecologically suitable areas, but where people are unwilling to coexist with elephants, i.e. areas where human-wildlife conflict may be exacerbated. This demonstrates opportunity for increasing sustainable coexistence potential by targeted conservation action in areas covering a significant proportion of our landscape, with suitable socio-ecological conditions but where people are unwilling to coexist with elephants.

## 5. Conclusion

We believe there is an urgent need to move beyond habitat suitability assessments for species based on their ecological requirements alone, and embrace the concept of holistic habitat suitability and sustainable coexistence potential. For such an endeavour, we argue and demonstrate that it is necessary to include spatially explicit willingness-to-coexist proxies in determining durable suitability of an area for wildlife. If local attitude towards wildlife and willingness to coexist cannot be quantified, perceived wildlife-induced threat reported by people could potentially serve as a valuable indicator of willingness to coexist.

When we asked respondents what they considered necessary to achieving sustainable coexistence, they repeatedly mentioned compensation, benefits and protection for both wildlife and people, as *“authorities concerned should also feel the loss I get”, “the solution is for [the] government to share the benefit with local people and employment to local people [is] required so the benefits can be seen by locals”* and *“there should be adequate protection for both people and wildlife”*. This latter point was also mentioned in concerns about the prioritisation of those organisations advocating on behalf of wildlife in the areas, a suggestion was made to be *“treating people as more important than elephants to reduce conflict*”.

The areas we highlight that are low in sustainable coexistence potential because people are not willing to coexist with elephants, suggest that if conservationists address reasons underlying the lack of willingness to coexist, persistence of wildlife species such as elephants could be better achieved, and sustainable coexistence realized. Only then can sustainable coexistence be realized. At the same time, strategic land use planning taking into consideration those areas with high willingness-to-coexist but low habitat suitability scores could increase the currently underused coexistence potential of these areas. Where feasible and realistic, this may necessitate active restoration of wildlife habitat in areas that are currently less suitable for them.

Finally, the looming threat posed by climatic change to humans, vegetation and wildlife (Bedelian and Ogutu, 2017; Hunninck et al., 2020), increases the urgency and importance of quantifying potentials of sustainable human-wildlife co-existence in ecosystems such as the Mara. Incorporating livelihood vulnerability and changes to habitat suitability linked to climatic and demographic changes is therefore an important next step in advancing our ability to reliably assess sustainable coexistence potential.

## Notes

### Competing Interest Statement

The authors have declared no competing interest.

