## SupplementaryInformation for "Identifying the potential for sustainable human–wildlife coexistence by integrating willingness to coexist with habitat suitability models"

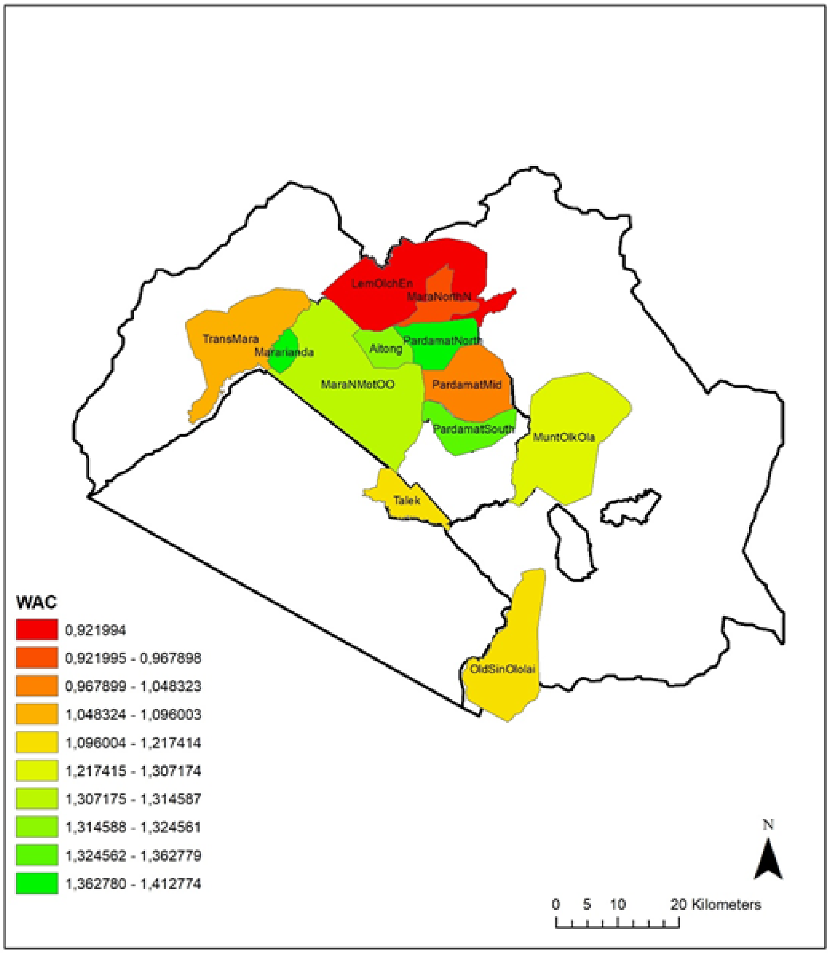


**Figure S1**. Distribution of the Wildlife Acceptance Capacity based on elephant density preference derived from bean simulations. Note that the answers have not been corrected for any reporting errors.

**Table S1.** Quantiles for willingness to coexist, average uncorrected Likert scale, and Wildlife Acceptance Capacity fall in.

| Cluster | Willingness to coexist | Uncorrected Likert | WAC |
|---|---|---|---|
| PardamatSouth | 1 | 2 | 1 |
| MuntOlkOla | 1 | 4 | 2 |
| MaraNorthN | 1 | 3 | 4 |
| PardamatNorth | 2 | 1 | 1 |
| Aitong | 2 | 2 | 2 |
| LemOlchEn | 2 | 2 | 4 |
| OldSinOlolai | 3 | 1 | 3 |
| TransMara | 3 | 4 | 3 |
| PardamatMid | 3 | 1 | 4 |
| Mararianda | 4 | 3 | 1 |
| MaraNMotOO | 4 | 3 | 2 |
| Talek | 4 | 4 | 3 |

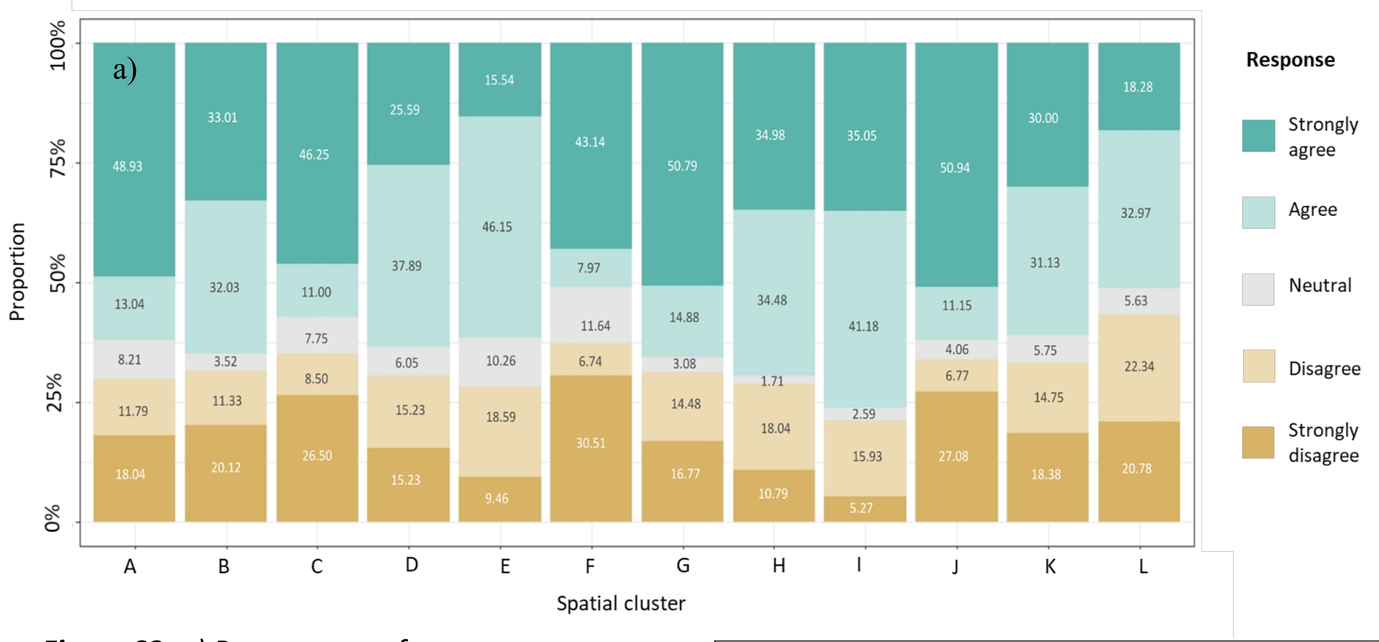


***Figure S2. a**) Percentages of responses to attitudinal statements, along a 5-point Likert scale ranging from 1) Strongly agree, 2) Agree, 3) Neutral, 4) Disagree, 5) Strongly disagree, for each of the spatial clusters (A=Greater Aitong area; B=Lemek, Olchoro Oiroua, Enonkishu; C=Mara North, Motorogi, Olare Orok conservancies; D=northern part of Mara North; E=Mara Rianta village; F=Proposed Muntoroben conservancy, Ol Kinyei, Olarro; G=Olderkesi Conservancy, Siana, Ololaimutiek village; H=central part of Paradamat; I= northern part of Paradamat; J=southern part of Paradamat; K=Talek town; L=Nyakweri Forest Conservation Area, Oloisukut, Masaai Moran). Negative statements were inverted to be positive for easy comparison across statements. **b)** Spatial clusters corresponding to the letters in figure a.*

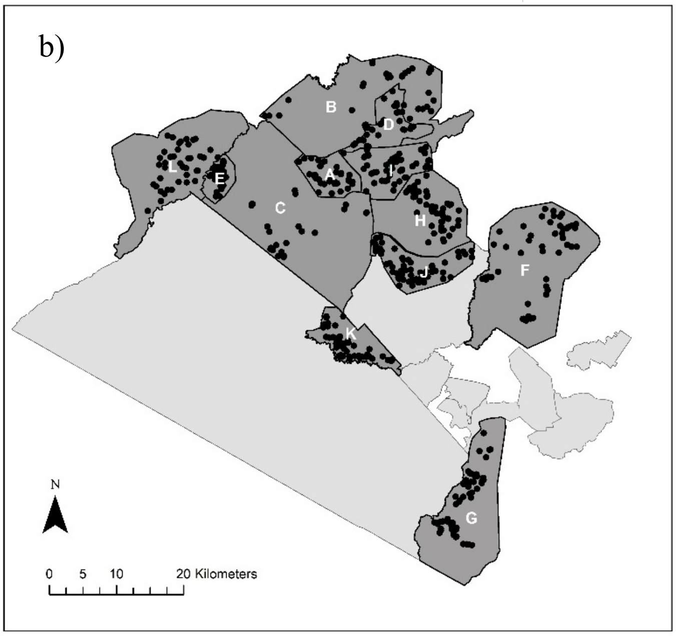

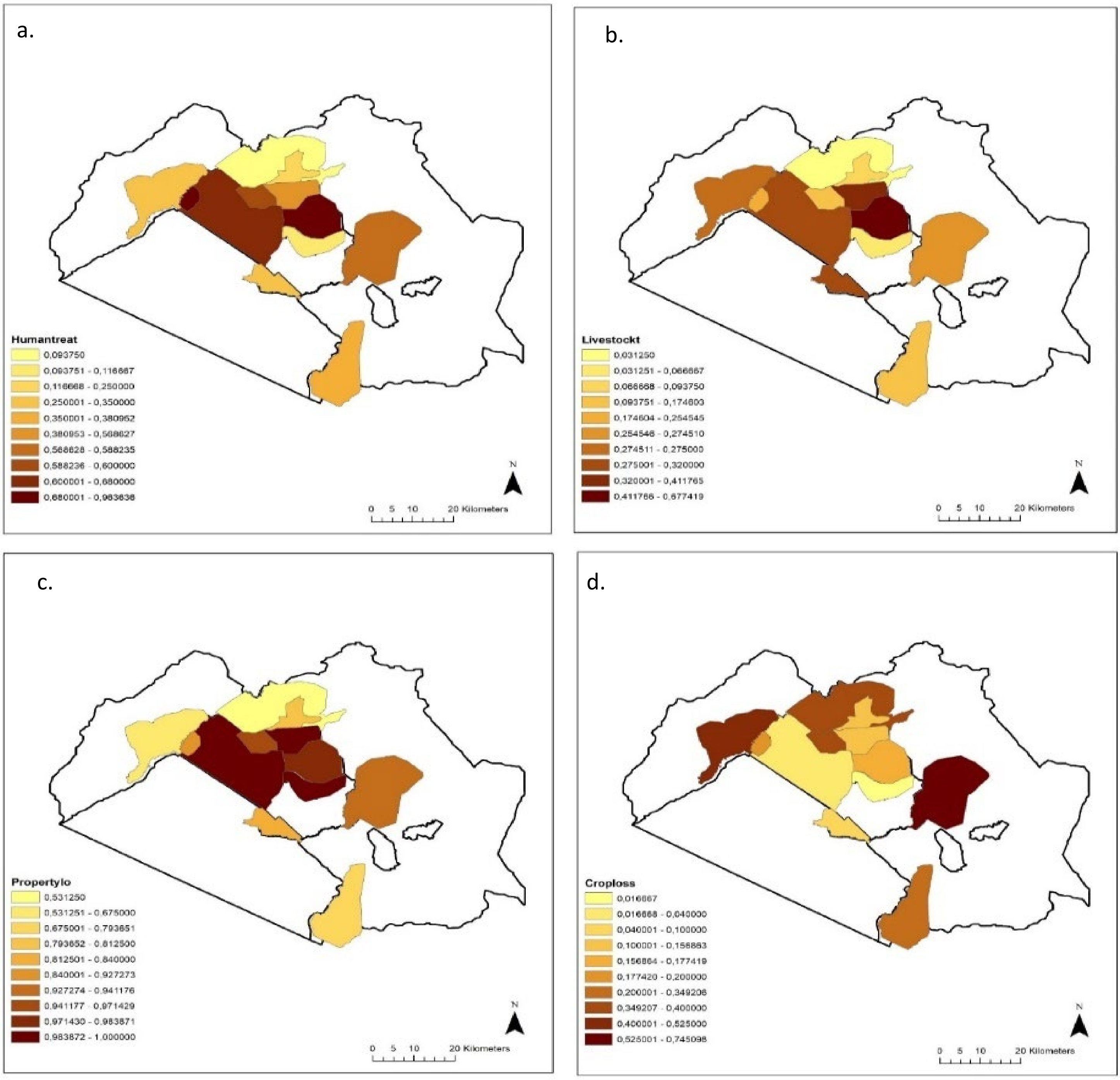


**Figure S3.** Distribution of reported issues people have coexisting with elephants across the Maasai Mara conservancies, for a) elephants as a threat to humans; b) elephants as threat to livestock; c) property loss due to elephants; and d) crop loss due to elephants.

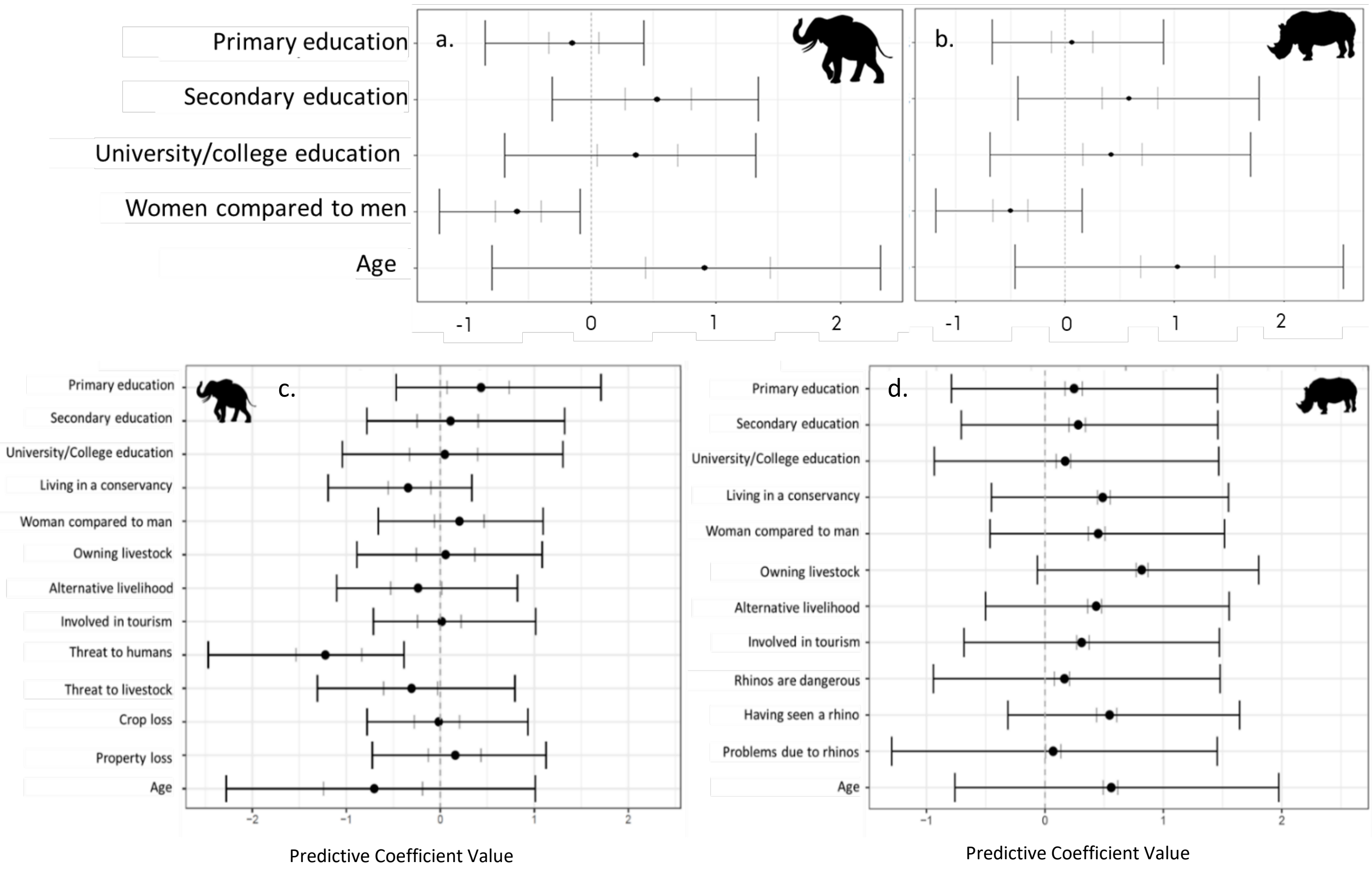


***Figure S4. a-b*** *Influence of covariates on the notional attitudes (φ) towards elephants (a) and rhinos (b) of n=556 respondents interviewed across the Maasai Mara conservancies, with 50% Credible Intervals (inner grey bars) and 95% Credible Intervals (outer bars), with the x-axes representing the coefficient values for each of the covariate.*

***c-d*** *Mean and variance of posterior predictive parameter estimates of willingness-to-coexist with elephants (c) and rhinos (d) across the Maasai Mara, of n=556 respondents interviewed across the Maasai Mara conservancies, with 50% Credible Intervals (inner grey bars) and 95% Credible Intervals (outer bars).*

**Table S2.** Parameter estimates for certain (c) and uncertain (uc) questions and their Credible Intervals, please see Supplementary Information 3.1.5 for detailed explanation of the Covariates.

| Species | Covariate | Mean | Q2.5 | Q25 | Q75 | Q97.5 |
|---|---|---|---|---|---|---|
| **Elephant** | $p_c^{11}$ | 1 | | | | |
| | $p_c^{10} = 1 - p_c^{11}$ | 0 | | | | |
| | $p_c^{01}$ | 0.626606 | 0.25658 | 0.574221 | 0.713435 | 0.769009 |
| | $p_{uc}^{11}$ | 0.650572 | 0.635934 | 0.644608 | 0.655684 | 0.669154 |
| | $p_{uc}^{10} = 1 - p_{uc}^{11}$ | 0.349428 | 0.364066 | 0.355392 | 0.344316 | 0.330846 |
| | $p_{uc}^{01}$ | 0.580775 | 0.502963 | 0.567366 | 0.601437 | 0.618961 |
| | $q_c^{11}$ | 0.611336 | 0.352286 | 0.487633 | 0.706184 | 0.876999 |
| | $q_c^{10} = 1 - q_c^{11}$ | 0.388664 | 0.647714 | 0.512367 | 0.293816 | 0.123001 |
| | $q_c^{01}$ | 0 | | | | |
| | $q_{uc}^{11}$ | 0.580158 | 0.538786 | 0.564383 | 0.595053 | 0.624103 |
| | $q_{uc}^{10} = 1 - q_{uc}^{11}$ | 0.419842 | 0.461214 | 0.435617 | 0.404947 | 0.375897 |
| | $q_{uc}^{01}$ | 0.550372 | 0.539431 | 0.546505 | 0.55415 | 0.56141 |
| **Rhino** | $p_c^{11}$ | 0.68777 | 0.661251 | 0.678777 | 0.696532 | 0.712794 |
| | $p_c^{10} = 1 - p_c^{11}$ | 0.31223 | 0.338749 | 0.321223 | 0.303468 | 0.287206 |
| | $p_c^{01}$ | 0 | | | | |
| | $p_{uc}^{11}$ | 0.676978 | 0.667976 | 0.673834 | 0.680073 | 0.686032 |
| | $p_{uc}^{10} = 1 - p_{uc}^{11}$ | 0.323022 | 0.332024 | 0.326166 | 0.319927 | 0.313968 |
| | $p_{uc}^{01}$ | 0.299947 | 0.177903 | 0.266015 | 0.341219 | 0.428718 |
| | $q_c^{11}$ | 0.949513 | 0.917659 | 0.939079 | 0.957315 | 0.971802 |
| | $q_c^{10} = 1 - q_c^{11}$ | 0.050487 | 0.082341 | 0.060921 | 0.042685 | 0.028198 |
| | $q_c^{01}$ | 0 | | | | |
| | $q_{uc}^{11}$ | 0.589967 | 0.579291 | 0.586249 | 0.593725 | 0.600747 |
| | $q_{uc}^{10} = 1 - q_{uc}^{11}$ | 0.410033 | 0.420709 | 0.413751 | 0.406275 | 0.399253 |
| | $q_{uc}^{01}$ | 0.426096 | 0.305364 | 0.384404 | 0.468593 | 0.554121 |

**Table S3**. Willingness to coexist for each of the covariate combinations for each of the spatial clusters. The combination column indicates the combination of socio-demographic characteristics of the group of people the specific willingness coexistence score is predicted for, written in the following order: 1) man or woman, 2) none, primary (prim), secondary (sec) or university (uni) formal education, 3) owning livestock or not owning livestock (no livestock), 4) alternative livelihood, or not having an alternative livelihood (no alternative). The cluster names are the following: EC=Enonkishu conservancy, LC= Lemek conservancy, MC=Motorogi conservancy, MNC=Mara north conservancy and Olare Orok conservancy, NC= Naboisho conservancy, ODC=Olderkesi conservancy, OKC=Ol Kinyei conservancy, OOC=Olchorro Oiroua conservancy, OTC=Oloisukut conservancy, OV=Ololaimutiek village, PCCA=proposed Pardamat community conservation area, PMCA=proposed Muntoroben conservancy area, PNEC=Proposed Naishi Enkutoto conservancy, POCA=Proposed Olderkesi conservancy area, SC=Siana conservancy, TA=Talek area.

| | Combination | EC | LC | MC | MNC | NC | ODC | OKC | OOC | OTC | OV | PCCA | PMCA | PNEC | POCA | PTM | SC | TA |
|---|---|---|---|---|---|---|---|---|---|---|---|---|---|---|---|---|---|---|
| 1 | man_none_nolivestock_noalternative | 0.465313 | 0.433841 | 0.34606 | 0.332245 | 0.752971 | 0.320663 | 0.584627 | 0.628 | 0.509289 | 0.429915 | 0.651235 | 0.734643 | 0.411491 | 0.444454 | 0.471411 | 0.447096 | 0.234847 |
| 2 | man_prim_nolivestock_noalternative | 0.569909 | 0.538488 | 0.446221 | 0.431044 | 0.822733 | 0.418174 | 0.681843 | 0.719927 | 0.612448 | 0.534509 | 0.7398 | 0.808263 | 0.515657 | 0.549178 | 0.575902 | 0.551824 | 0.318497 |
| 3 | man_sec_nolivestock_noalternative | 0.442127 | 0.411018 | 0.325202 | 0.311823 | 0.73516 | 0.300632 | 0.561743 | 0.605894 | 0.485904 | 0.40715 | 0.629696 | 0.716009 | 0.389036 | 0.421489 | 0.448176 | 0.424098 | 0.218453 |
| 4 | man_uni_nolivestock_noalternative | 0.426789 | 0.395993 | 0.311654 | 0.298586 | 0.722827 | 0.287671 | 0.546318 | 0.590892 | 0.470327 | 0.392172 | 0.615024 | 0.703144 | 0.374304 | 0.406345 | 0.432791 | 0.408927 | 0.207982 |
| 5 | man_none_livestock_noalternative | 0.3996 | 0.369501 | 0.288113 | 0.275637 | 0.699803 | 0.265244 | 0.518401 | 0.563526 | 0.442506 | 0.365781 | 0.588147 | 0.679211 | 0.348426 | 0.379596 | 0.40549 | 0.382117 | 0.190109 |
| 6 | man_prim_livestock_noalternative | 0.50333 | 0.471555 | 0.381282 | 0.36685 | 0.780197 | 0.354702 | 0.62107 | 0.662832 | 0.547222 | 0.467569 | 0.684983 | 0.763254 | 0.448802 | 0.482305 | 0.509452 | 0.484975 | 0.263307 |
| 7 | man_sec_livestock_noalternative | 0.377377 | 0.347983 | 0.26931 | 0.257354 | 0.679788 | 0.247415 | 0.495019 | 0.540393 | 0.419565 | 0.344361 | 0.565312 | 0.658493 | 0.327497 | 0.357823 | 0.383149 | 0.360284 | 0.176119 |
| 8 | man_uni_livestock_noalternative | 0.362824 | 0.333955 | 0.257203 | 0.245603 | 0.666049 | 0.235974 | 0.479423 | 0.524852 | 0.404441 | 0.330405 | 0.549913 | 0.644318 | 0.313898 | 0.343608 | 0.368505 | 0.346023 | 0.167243 |

| | | | | | | | | | | | | | | | | | | |
|---|---|---|---|---|---|---|---|---|---|---|---|---|---|---|---|---|---|---|
| 9 | man_none_nolivestock_alternative | 0.439722 | 0.408659 | 0.323065 | 0.309734 | 0.733256 | 0.298585 | 0.559339 | 0.603562 | 0.483467 | 0.404797 | 0.627418 | 0.714021 | 0.38672 | 0.419112 | 0.445765 | 0.421717 | 0.216792 |
| 10 | man_prim_nolivestock_alternative | 0.544423 | 0.512732 | 0.420854 | 0.405907 | 0.80716 | 0.393269 | 0.659021 | 0.69863 | 0.587659 | 0.508734 | 0.719424 | 0.79174 | 0.489834 | 0.523491 | 0.550491 | 0.526157 | 0.296503 |
| 11 | man_sec_nolivestock_alternative | 0.416817 | 0.386257 | 0.302951 | 0.290094 | 0.714562 | 0.279366 | 0.536166 | 0.580972 | 0.460155 | 0.38247 | 0.605299 | 0.69454 | 0.364778 | 0.396521 | 0.422783 | 0.399082 | 0.201327 |
| 12 | man_uni_nolivestock_alternative | 0.401725 | 0.371565 | 0.289932 | 0.277407 | 0.701659 | 0.266972 | 0.52061 | 0.565701 | 0.44469 | 0.367837 | 0.590289 | 0.681136 | 0.350438 | 0.381682 | 0.407625 | 0.384209 | 0.191475 |
| 13 | man_none_livestock_alternative | 0.375088 | 0.345772 | 0.267394 | 0.255493 | 0.677661 | 0.245602 | 0.492581 | 0.537969 | 0.417191 | 0.342162 | 0.562913 | 0.656296 | 0.325352 | 0.355584 | 0.380846 | 0.358038 | 0.174708 |
| 14 | man_prim_livestock_alternative | 0.477516 | 0.445907 | 0.357225 | 0.343199 | 0.761968 | 0.331424 | 0.596469 | 0.639368 | 0.521521 | 0.441957 | 0.662275 | 0.74408 | 0.423401 | 0.456577 | 0.483629 | 0.459232 | 0.243762 |
| 15 | man_sec_livestock_alternative | 0.353426 | 0.324924 | 0.249469 | 0.238107 | 0.656894 | 0.228682 | 0.469228 | 0.514647 | 0.394633 | 0.321423 | 0.539774 | 0.634894 | 0.30516 | 0.334446 | 0.359043 | 0.33683 | 0.161626 |
| 16 | man_uni_livestock_alternative | 0.339293 | 0.311382 | 0.237964 | 0.226967 | 0.642688 | 0.217857 | 0.453714 | 0.499043 | 0.37982 | 0.307961 | 0.524231 | 0.620304 | 0.292086 | 0.320696 | 0.344805 | 0.32303 | 0.153344 |
| 17 | woman_none_nolivestock_noalternative | 0.542191 | 0.510484 | 0.418663 | 0.403739 | 0.805755 | 0.391124 | 0.656996 | 0.696732 | 0.585477 | 0.506485 | 0.717605 | 0.790253 | 0.487586 | 0.521246 | 0.548264 | 0.523914 | 0.29463 |
| 18 | woman_prim_nolivestock_noalternative | 0.643278 | 0.613582 | 0.523031 | 0.507636 | 0.863317 | 0.494466 | 0.744671 | 0.777687 | 0.682601 | 0.609781 | 0.79463 | 0.851562 | 0.591649 | 0.623748 | 0.648878 | 0.626254 | 0.388755 |
| 19 | woman_sec_nolivestock_noalternative | 0.518892 | 0.487098 | 0.39608 | 0.381433 | 0.790691 | 0.369084 | 0.635613 | 0.676607 | 0.562604 | 0.483101 | 0.698265 | 0.774323 | 0.464254 | 0.497868 | 0.525003 | 0.500541 | 0.275566 |
| 20 | woman_uni_nolivestock_noalternative | 0.503293 | 0.471518 | 0.381247 | 0.366816 | 0.780172 | 0.354668 | 0.621035 | 0.662798 | 0.547186 | 0.467532 | 0.684951 | 0.763227 | 0.448766 | 0.482268 | 0.509415 | 0.484938 | 0.263279 |
| 21 | woman_none_livestock_noalternative | 0.475271 | 0.443685 | 0.355162 | 0.341173 | 0.760332 | 0.329433 | 0.594301 | 0.63729 | 0.519275 | 0.439739 | 0.660259 | 0.742363 | 0.421205 | 0.454346 | 0.481383 | 0.456998 | 0.242107 |
| 22 | woman_prim_livestock_noalternative | 0.57968 | 0.548407 | 0.45612 | 0.440874 | 0.828489 | 0.427931 | 0.690451 | 0.727916 | 0.621893 | 0.544441 | 0.747421 | 0.814382 | 0.525634 | 0.559055 | 0.585635 | 0.561689 | 0.327237 |
| 23 | woman_sec_livestock_noalternative | 0.452009 | 0.420729 | 0.334036 | 0.320466 | 0.74287 | 0.309104 | 0.571559 | 0.615398 | 0.495894 | 0.416834 | 0.638968 | 0.724067 | 0.398579 | 0.431266 | 0.458082 | 0.433891 | 0.225355 |
| 24 | woman_uni_livestock_noalternative | 0.436596 | 0.405593 | 0.320294 | 0.307025 | 0.730765 | 0.295932 | 0.556207 | 0.600519 | 0.480296 | 0.401741 | 0.624444 | 0.711421 | 0.383712 | 0.416023 | 0.442629 | 0.418623 | 0.214644 |
| 25 | woman_none_nolivestock_alternative | 0.516456 | 0.484661 | 0.393748 | 0.379134 | 0.789072 | 0.366815 | 0.633351 | 0.674469 | 0.560202 | 0.480665 | 0.696205 | 0.772614 | 0.461828 | 0.495429 | 0.52257 | 0.498102 | 0.273622 |
| 26 | woman_prim_nolivestock_alternative | 0.619235 | 0.588817 | 0.497217 | 0.481815 | 0.850661 | 0.468678 | 0.724536 | 0.759313 | 0.659807 | 0.584937 | 0.777256 | 0.838022 | 0.566472 | 0.599209 | 0.624992 | 0.601774 | 0.364504 |
| 27 | woman_sec_nolivestock_alternative | 0.493072 | 0.461343 | 0.37165 | 0.357372 | 0.773079 | 0.345367 | 0.611366 | 0.653601 | 0.537037 | 0.457369 | 0.676062 | 0.755759 | 0.438674 | 0.472068 | 0.499194 | 0.474733 | 0.255425 |

| | | | | | | | | | | | | | | | | | |
|---|---|---|---|---|---|---|---|---|---|---|---|---|---|---|---|---|---|
| 28 | woman_uni_nolivestock_alternative | 0.477479 | 0.445871 | 0.357191 | 0.343165 | 0.761941 | 0.331391 | 0.596433 | 0.639333 | 0.521484 | 0.44192 | 0.662241 | 0.744052 | 0.423364 | 0.45654 | 0.483592 | 0.459195 | 0.243734 |
| 29 | woman_none_livestock_alternative | 0.449593 | 0.418353 | 0.331869 | 0.318345 | 0.741002 | 0.307024 | 0.569168 | 0.613087 | 0.493455 | 0.414465 | 0.636714 | 0.722113 | 0.396243 | 0.428874 | 0.455661 | 0.431496 | 0.223657 |
| 30 | woman_prim_livestock_alternative | 0.55432 | 0.522714 | 0.430628 | 0.415582 | 0.813306 | 0.402848 | 0.667946 | 0.706979 | 0.597311 | 0.51872 | 0.727423 | 0.798255 | 0.499827 | 0.533452 | 0.560362 | 0.536112 | 0.304909 |
| 31 | woman_sec_livestock_alternative | 0.426566 | 0.395776 | 0.311459 | 0.298396 | 0.722645 | 0.287485 | 0.546093 | 0.590672 | 0.4701 | 0.391955 | 0.614808 | 0.702955 | 0.374091 | 0.406125 | 0.432567 | 0.408707 | 0.207832 |
| 32 | woman_uni_livestock_alternative | 0.411369 | 0.380947 | 0.29823 | 0.285492 | 0.709959 | 0.274868 | 0.530578 | 0.575496 | 0.454582 | 0.377181 | 0.599921 | 0.689755 | 0.359591 | 0.39116 | 0.417313 | 0.39371 | 0.197741 |

### Model evaulations

#### Model evaluation SDMs

Evaluating the goodness of fit of ZANB hurdle SDMs is challenging because the models consist of two parts, with (a) a binomial model of the probability of zeros—for unsuitable sites, followed by (b) a truncated negative binomial model for non-zero observations—for the distribution of elephant density in suitable sites. Therefore, we evaluated model fit using diagnostic plots, including hanging Rootograms, residual QQ plots, plots of observed versus expected values and residual versus fitted values (Kleiber, 2016; Wang, 2020).

All models under fitted larger elephant densities, as these values were also rare in the observed data. Based on plots of the observed versus predicted variables and residuals versus fitted values, the nested model containing only the covariates with high predictive power outperformed the full model containing all the selected ecological and social variables. However, both models performed well on the hanging Rootograms and residual QQ plots. As a result, we selected the model containing, as covariates: water, NDVI, elevation, total number of settlements and precipitation for further analyses (Kleiber, 2016; Wang, 2020).

#### Model evaluation INLA

Following Blangiardo (2013), we used several methods that take into account the complex nuances of approximate Bayesian models and the effect of zero-inflation to evaluate model fit, namely the Conditional Predictive Ordinate (CPO), Probability Integral Transform (PIT), failure scores and posterior predictive p-value distributions. The smaller the AIC and failure scores or the larger the CPO scores the better is the model fit. A histogram of PIT measure that tends towards uniformity also indicates a good model fit. Moreover, we interpreted, a scatterplot of the posterior mean of the predictive distribution against the observed values, as reflecting an average prediction close to the observations, and a histogram of the posterior predictive p-value without strong variation in the distribution as indicative of a good model fit (Blangiardo, 2013).

While the AIC and the CPO for the other models suggested a slightly better fit, PIT and posterior predictive checks revealed the model that best fitted the dry season data included the covariates: water, elevation, NDVI, slope, livestock biomass and total number of houses. For the wet season, only the histogram of the PIT measure for the model containing only the ecological variables: water, elevation, NDVI and slope tended towards uniformity, yet the AIC scores revealed strong model fit issues compared to the other models.
